# GFAP-directed Inactivation of *Men1* Exploits Glial Cell Plasticity in Favor of Neuroendocrine Reprogramming

**DOI:** 10.1101/2022.02.10.479845

**Authors:** Suzann Duan, Travis W. Sawyer, Ricky A. Sontz, Bradley A. Wieland, Andres F. Diaz, Juanita L. Merchant

## Abstract

**BACKGROUND & AIMS:** Efforts to characterize the signaling mechanisms that underlie gastroenteropancreatic neoplasms (GEP-NENs) are precluded by a lack of comprehensive model systems that recapitulate pathogenesis. Investigation into a potential cell-of-origin for gastrin-secreting NENs revealed a role for enteric glia in neuroendocrine cell specification. Here we investigated the hypothesis that loss of menin in glial cells stimulated neuroendocrine differentiation and tumorigenesis.

**METHODS:** Using *Cre-lox* technology, we generated a conditional glial fibrillary acidic protein-directed *Men1* knockout (*GFAP*^*ΔMen1*^) mouse model. *Cre* specificity was confirmed using a tdTomato reporter. *GFAP*^*ΔMen1*^ mice were evaluated for GEP-NEN development and neuroendocrine cell hyperplasia. siRNA-mediated *Men1* silencing in a rat enteric glial cell line was performed in parallel.

**RESULTS:** *GFAP*^*ΔMen1*^ mice developed pancreatic NENs, in addition to pituitary prolactinomas that phenocopied the human MEN1 syndrome. *GFAP*^*ΔMen1*^ mice exhibited gastric neuroendocrine hyperplasia that coincided with a significant loss of GFAP expression. Mechanistically, *Men1* deletion induced reprogramming from a mature glial phenotype toward a neuroendocrine lineage. Furthermore, blockade of Hedgehog signaling in enteric glia attenuated neuroendocrine hyperplasia by restricting the neuroendocrine cell fate.

**CONCLUSIONS:** GFAP-directed *Men1* inactivation exploits glial cell plasticity in favor of neuroendocrine differentiation.

**GRAPHICAL ABSTRACT:** 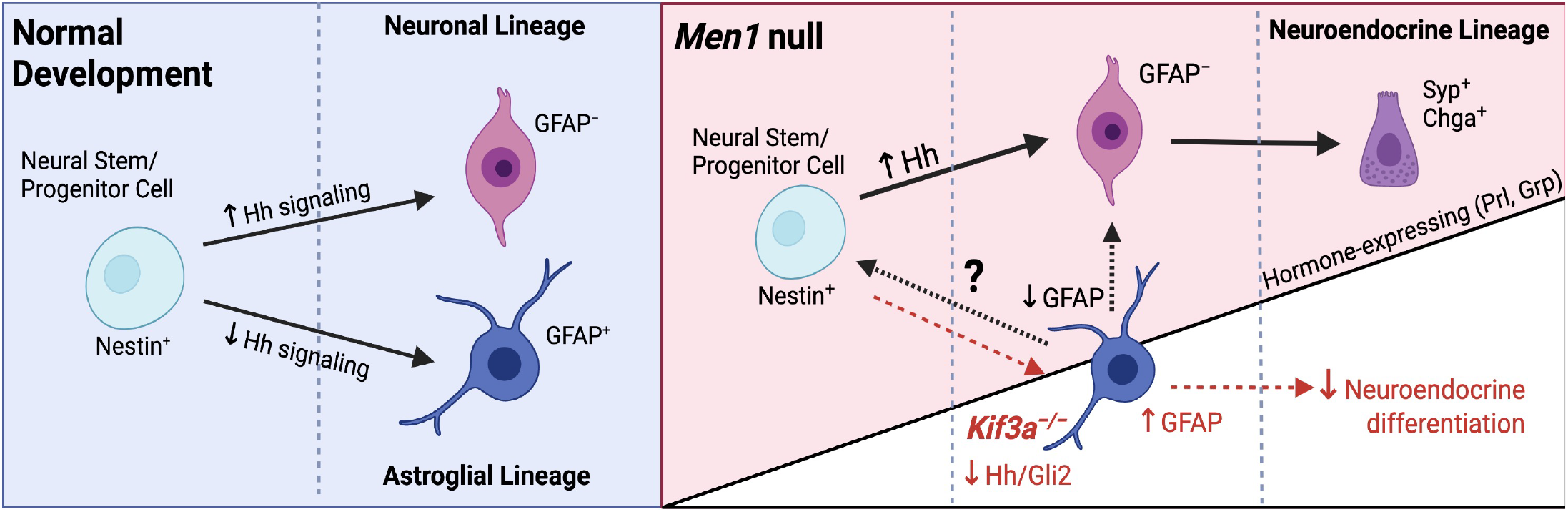

## INTRODUCTION

Gastroenteropancreatic neuroendocrine neoplasms (GEP-NENs) comprise a heterogeneous group of malignancies showing an increase in incidence and prevalence across the Unites States ^[1,2]^. GEP-NENs are comprised of endocrine-producing cells and include gastric carcinoids, gastrinomas, and pancreatic neuroendocrine tumors (NETs) ^[3]^. The development of GEP-NENs is frequently associated with sporadic and inherited mutations in the Multiple Endocrine Neoplasia I (*MEN1*) gene ^[4]^. Inactivation of the *MEN1* locus causes loss of the tumor suppressor protein menin and coincides with the development of endocrine tumors in the pancreas, pituitary, and upper GI tract ^[5]^. Patients carrying a *MEN1* mutation are predisposed to developing gastrinomas, a GI NET that produces excess levels of gastrin, a peptide hormone that stimulates acid secretion, inflammation, and proliferation ^[6,7,8,9]^. Such *MEN1*-associated gastrinomas preferentially develop in Brunner’s glands located within the duodenal submucosa, with an estimated >50% of MEN1 gastrinomas exhibiting lymph node metastases at the time of diagnosis ^[10,11]^.

Loss of menin function is a critical event underlying the formation of *MEN1* gastrinomas however, the signaling cues that regulate menin-mediated suppression of gastrin remain elusive. Homozygous deletion of *Men1 in utero* is embryonically lethal in mice, whereas heterozygous inactivation promotes endocrine tumors of the pancreas and pituitary, but not in the GI tract ^[12]^. Importantly, these studies did not identify potential cells-of-origin for tumor development. Subsequently, we reported the development of the first genetically engineered mouse model that displays gastric carcinoids ^[13]^. Conditional deletion of *Men1* from the GI tract epithelium using the *Villin Cre* transgene on a somatostatin-null genetic background resulted in antral G cell hyperplasia, hypergastrinemia, and gastric carcinoids ^[13]^. Systemic gastric acid suppression using a proton pump inhibitor accelerated carcinoid development and coincided with the emergence of hyperplastic gastrin-expressing glial cells within the lamina propria of the duodenum ^[14]^. The plasticity of glial cells directed towards an endocrine phenotype coincided with decreased menin ^[14]^. Taken together, these observations suggest that hyperplastic G-cells might emerge from reprogrammed neural crest-derived cells rather than from endoderm-derived enteroendocrine cells. Indeed, prior studies have shown that multipotent glial cells can generate neuroendocrine cells during normal development ^[15]^ and upon overexpression of oncogenes such as *MYCN* ^[16]^. Therefore, we tested the hypothesis that loss of menin in enteric glial cells promotes neuroendocrine cell differentiation. Here we show that conditional deletion of *Men1* in GFAP-expressing glial cells results in NEN development in the pituitary and pancreas, in addition to neuroendocrine hyperplasia in the stomach. These events coincided with loss of the glial phenotype and emergence of an endocrine phenotype with tissue-specific hormone expression, suggesting a direct role for glial cell reprogramming upon removal of menin.

## RESULTS

### *GFAP*^*ΔMen1*^ mice develop pancreatic neuroendocrine neoplasms (PNENs)

To determine whether loss of menin in glial cells was sufficient to drive the development of GEP-NENs, we used *Cre-lox* technology to conditionally delete *Men1* under the control of the <u>g</u>lial <u>f</u>ibrillary <u>a</u>cidic <u>p</u>rotein (GFAP) promoter (*GFAP*^*ΔMen1*^). By 18–24 months of age, 7 of 12 *GFAP*^*ΔMen1*^ mice developed macroscopic lesions throughout the pancreas (Table 1). Histological analysis of the pancreas identified significant islet hyperplasia and features characteristic of neuroendocrine neoplasms (Figure 1A). Pancreatic lesions varied from poorly differentiated with a solid sheet-like morphology to highly differentiated with a rosette pattern (Figure 1B–D). Consistent with their unique histology, highly differentiated neoplasms exhibited cytoplasmic or absent menin expression (Figure 1E) and strong immunoreactivity for the neuroendocrine markers synaptophysin (Syp) and chromogranin A (ChgA), in addition to the alpha cell specification factor iroquois homeobox protein 2 (Irx2) (Figure 1E and F). Poorly differentiated tumors also exhibited strong nuclear expression of Irx2 with a sparse number of cells displaying immunoreactivity for Syp (Figure 1G). Based on the mitotic index used to categorize human pancreatic NENs ^[17]^, we stained *GFAP*^*ΔMen1*^ neoplasms for Ki-67 and classified the tumors as PNETs (<2% per high power field, HPF) (Figure 1H and I) or PNECs (>20% per HPF) (Figure 1J). Further characterization of highly differentiated PNETs revealed the presence of both hyperplastic insulin-expressing beta-cells and glucagon-expressing alpha-cells (Figure 1K–M). Evaluation of neuroendocrine transcript expression revealed a 10-fold increase in *ChgA* and 5-fold increase in *Syp* mRNA, however no significant differences in hormone expression were observed between wild type pancreas and the PNETs (Figure 1N). Next, we evaluated whether *GFAP*^*ΔMen1*^ PNENs express GFAP. As previously reported, GFAP expression were primarily localized to the periphery of islets in both wild type and *GFAP*^*ΔMen1*^ mice ^[18]^, however expression was uniquely lost surrounding PNETs (Figure 1O). In comparison, PNECs exhibited robust GFAP expression. Consistent with this, the expression of glial transcripts *Gfap* and *S100b* were significantly lower in PNETs compared to the pancreas of wild type mice, while *Men1* mRNA exhibited a trending decrease (Figure 1P). In summary, development of well-differentiated PNETs in this model coincided with loss of GFAP expression from the periphery of normal endocrine cells.

**Table 1.** Number of mice with macroscopic phenotypes and age/sex-dependent differences

| **Men1** | **Pancreatic**<br>**hyperplasia/tumor** | **Pituitary**<br>**Adenoma** | **Antral**<br>**hyperplasia/tumor** | **BG/Duodenal**<br>**Polyp** |
| --- | --- | --- | --- | --- |
| WT | 0/20 Avg. age 17.5 mo | 0/20 | 0/20 | 0/20 |
| FL/+ | 1/9 Avg. age 17.9 mo | 1/9. Male<br>1/7 Female | 1/9 | 1/9 |
| FL/FL | 7/26 Avg. age 17.9 mo<br>**7/12** 18mo and older<br>0/14 Under 18mo | 15/29<br>**15/22** Female<br>1/7 Male | 8/26 | 2/26 |

**Figure 1.**
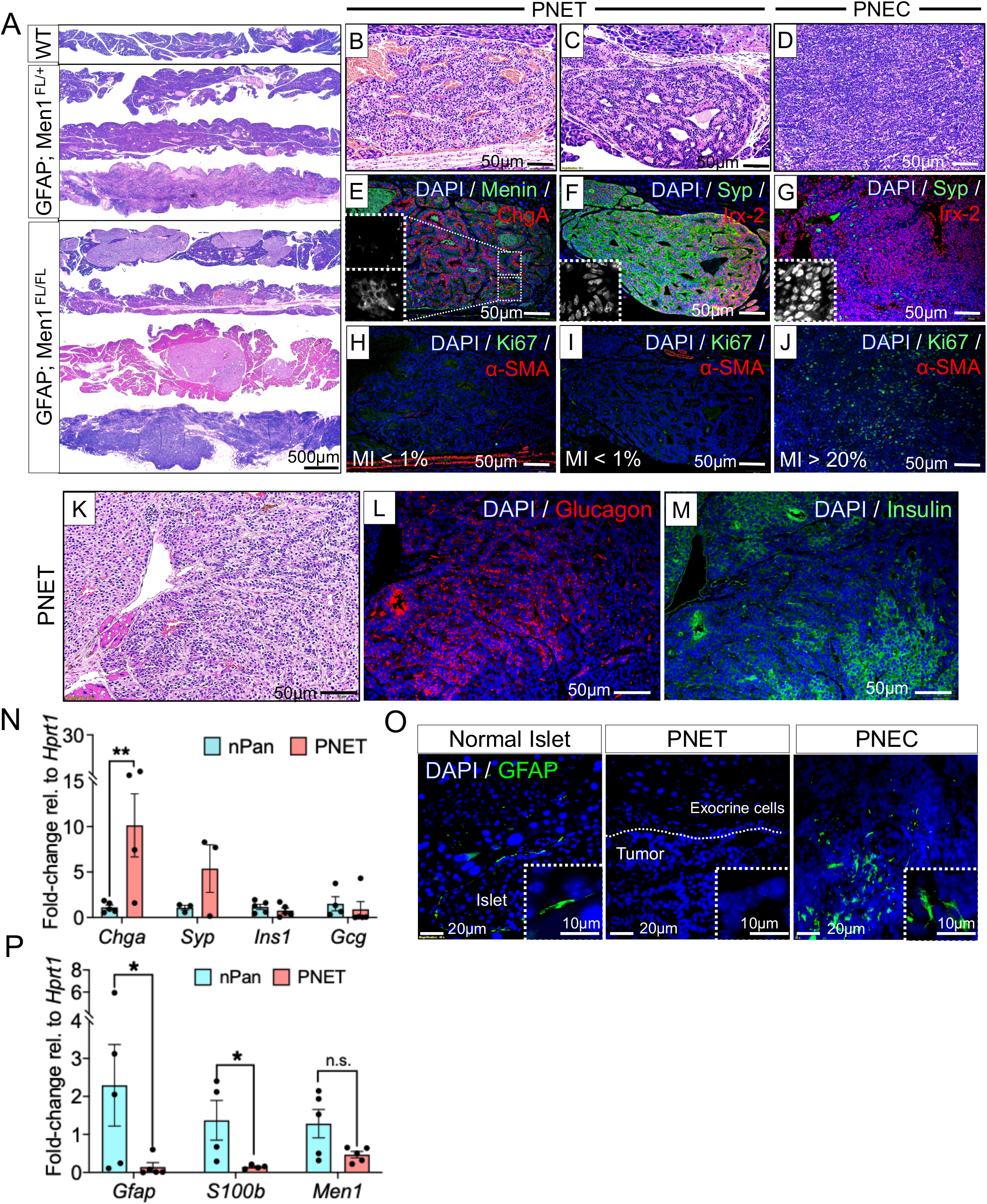
GFAP-directed inactivation of *Men1* promotes pancreatic islet hyperplasia and the development of pancreatic neuroendocrine neoplasms. (A) H&Es of well differentiated and poorly differentiated tumors in *GFAP*^*△Men1*^ mice compared to wild type (WT) and heterozygous groups. (B) and (C) H&Es of well differentiated *GFAP*^*△Men1*^ PNETs compared to (D) a poorly differentiated PNEC (E) Immunofluorescent staining of a PNET for chromogranin A and menin. Insets: tumor areas with absent and cytoplasmic menin (white). (F) Immunofluorescent staining of a PNET and (G) PNEC for synaptophysin and the alpha cell-specification factor Iriquois-2 (Irx-2). Insets: Irx-2 is expressed in the nucleus of tumor cells (white). (H) and (I) Ki-67 and smooth muscle actin (SMA) staining of PNETs compared to (J) a PNEC. (K) H&E of a *GFAP*^*△Men1*^ PNET stained for (L) glucagon and (M) insulin. (N) Quantitation of neuroendocrine-related transcripts in WT pancreas and *GFAP*^*△Men1*^ PNETs. (O) Immunofluorescent staining of *GFAP*^*△Men1*^ PNETs for GFAP compared to normal islets and a poorly differentiated PNEC. (P) Quantitation of *GFAP, S100b*, and *Men1* mRNA in WT pancreas and *GFAP*^*△Men1*^ PNETs. n=4–5 mice per group; * *p* < 0.05, ** *p* < 0.01 by unpaired Student’s T-test. Data are represented as mean ± SEM.

### *GFAP*^*ΔMen1*^ mice develop PitNETs that phenocopy human MEN1 prolactinomas

Following our observation of hydrocephalus-like neurological symptoms in a subset of *GFAP*^*ΔMen1*^ mice, we further investigated the brain for potential effects of GFAP-directed *Men1* inactivation. GFAP is the main intermediate filament protein in brain astrocytes and related cell types such as enteric glial cells where it is strongly expressed ^[19]^. Moreover, GFAP marks nestin^+^ subventricular neural stem cells ^[20–22]^ and interacts with menin ^[23,24]^. Fifty percent of *GFAP*^*ΔMen1*^ mice exhibited hemorrhagic pituitary tumors, with the phenotype being nearly exclusively sex-dependent. Female *GFAP*^*ΔMen1*^ mice (15 of 22 or 68%) developed macroscopic pituitary adenomas as early as 15 months of age (Table 1, Figure 2A and B). Subsequent immunostaining of *GFAP*^*ΔMen1*^ adenomas demonstrated diffuse immunoreactivity for synaptophysin and chromogranin A (Figure 2C). Strong staining for prolactin and the pituitary-specific marker Pit-1 showed these tumors to be prolactinomas (Figure 2D and E). Additionally, variable Ki-67 expression appeared to inversely correlate with the expression of neuroendocrine markers, with stronger neuroendocrine status associated with lower Ki-67 staining (Figure 2E). Further evaluation of GFAP and menin expression in *GFAP*^*ΔMen1*^ prolactinomas showed absent GFAP staining and low expression of menin within the tumor compared to the adjacent hypothalamus (Figure 2F and G). Similar to the previous observations in PNETs, *Gfap* and *Men1* mRNA expression was significantly reduced in prolactinomas compared to wild type pituitary (Figure 2H). As expected, the neuroendocrine transcripts *ChgB, Syp*, and *Enolase 2* (*Eno2*) were significantly elevated in *GFAP*^*ΔMen1*^ pitNETs compared to wild type pituitary. Additionally, mRNA levels of the *prolactin* and *vasoactive intestinal peptide* (*Vip*) hormones were 15-fold higher in pitNETs (Figure 2I). To determine whether GFAP-negative *GFAP*^*ΔMen1*^ pitNETs share features with neural stem cells, we generated tumor neurospheres from a *GFAP*^*ΔMen1*^ pituitary prolactinoma. Subsequent immunostaining of *GFAP*^*ΔMen1*^ tumor neurospheres revealed that tumor cells co-expressed prolactin and the neural stem cell markers Sox2 and Nestin, indicating that the hormone-expressing tumor cells arose from a neural stem cell lineage (Figure 2J).

**Figure 2.**
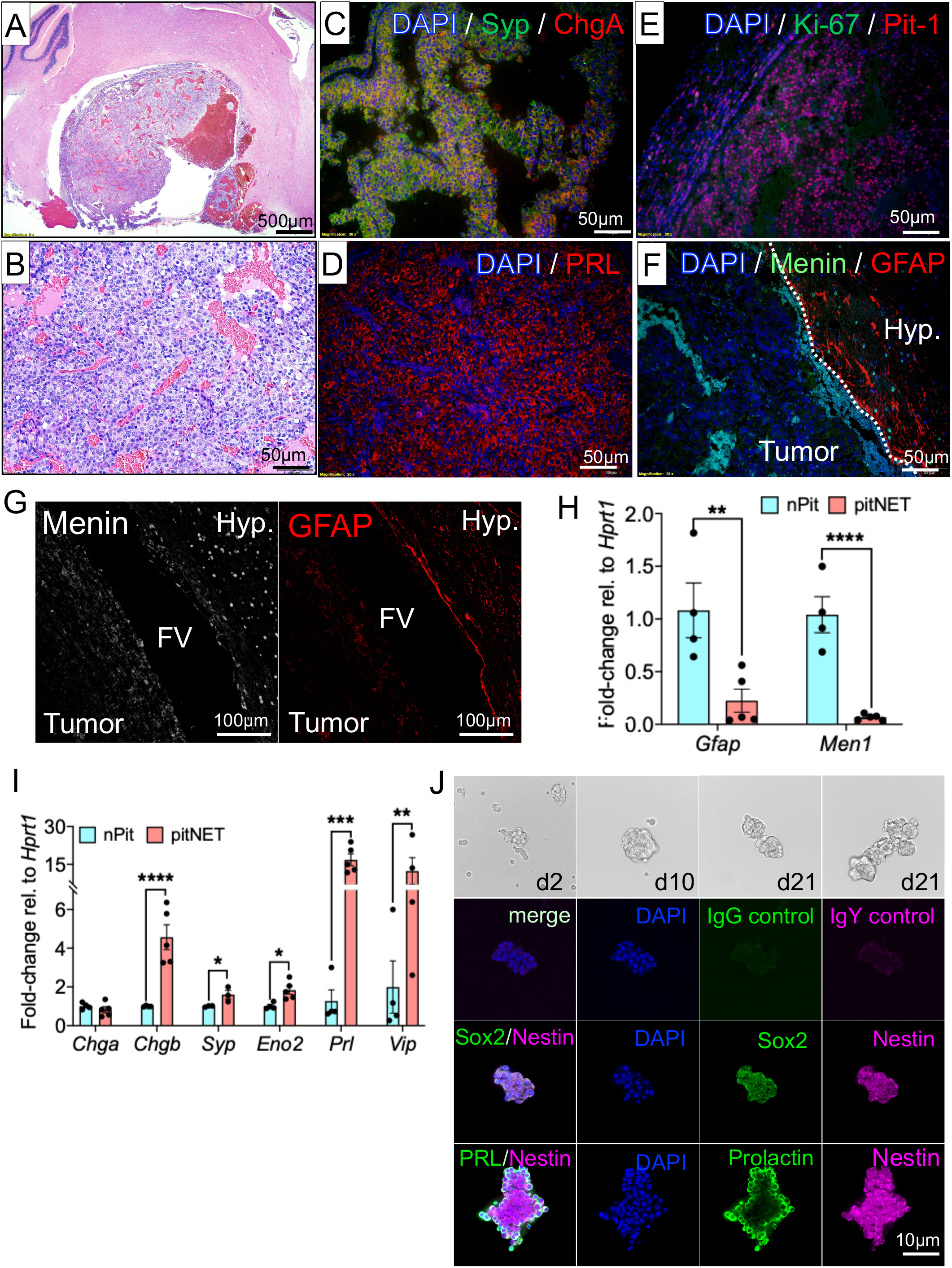
*GFAP*^*ΔMen1*^ mice develop pituitary neuroendocrine tumors (pitNETs) that phenocopy human MEN1 prolactinomas. (A) and (B) H&Es of a sagittal section of a *GFAP*^*ΔMen1*^ pitNET. (C) Immunofluorescent staining of a *GFAP*^*ΔMen1*^ pitNET for synaptophysin and chromogranin A, (D) prolactin and (E) Ki-67 and Pit-1. (F) Immunofluorescent staining of a *GFAP*^*ΔMen1*^ pitNET for GFAP and menin. (G) Tumors show low expression of menin and GFAP compared to the adjacent hypothalamus (Hyp). FV = fourth ventricle. (H) Quantitation of GFAP mRNA in WT pituitary and *GFAP*^*ΔMen1*^ pitNETs. (I) Quantitation of neuroendocrine-related transcripts in WT pituitary and *GFAP*^*ΔMen1*^ pitNETs. n=4–5 mice per group with the exception of wild type pituitary group which represents four samples of three pooled pituitaries for a total of 12 tissues in this group; * *p* < 0.05, ** *p* < 0.01, *** *p* < 0.001, **** *p* < 0.0001 by unpaired Student’s T-test. Data are represented as mean ± SEM. (J) Brightfield images of cultured tumor neurospheres derived from a *GFAP*^*ΔMen1*^ pitNET showing growth at days (d) 2, 10, and 21. Tumor neurospheres were stained for prolactin (PRL) and the neural stem cell markers Nestin and Sox2. IgG and IgY secondary antibodies served as negative controls.

### Hyperplastic reprogramming of the gastro-intestinal epithelium in *GFAP*^*ΔMen1*^ mice

To determine whether loss of menin in enteric glia supports the development of NENs in the luminal gastrointestinal (GI) tract, we evaluated the stomach and proximal small intestine of *GFAP*^*ΔMen1*^ mice for changes in enteroendocrine cell numbers and composition. Although no gastrinomas or small intestinal NENs were observed at 18–24 months of age, *GFAP*^*ΔMen1*^ mice showed hyperplastic changes in both the corpus and gastric antrum by nine months of age (Figure 3A). Further analysis of neuroendocrine marker expression revealed a ∼two-fold increase in *ChgA* transcripts in both the corpus and gastric antra of *GFAP*^*ΔMen1*^ mice, while no differences in mRNA levels were observed in the proximal duodenum (Figure 3B). Subsequent immunostaining confirmed increased *ChgA* expression in the proximal and distal stomach (Figure 3C). Moreover, the *GFAP*^*ΔMen1*^ mice exhibited a significant increase in antral *Syp* expression (Figure 3D). Elevated Syp protein in gastric antra was further confirmed by immunostaining (Figure 3E). As menin is a known repressor of gastrin gene expression and its loss in intestinal epithelium was previously shown to stimulate hypergastrinemia in *Villin Cre; Men1*^*Fl/FL*^ mice ^[13]^, we assessed whether deletion of menin in enteric glia similarly induced gastrin hormone expression. Consistent with the previous increases in neuroendocrine marker expression, we observed a significant increase (three-fold) in gastrin mRNA and gastrin immunostaining in the antra of *GFAP*^*ΔMen1*^ mice (Figure 3F and G). Therefore, conditional deletion of menin in enteric glial cells stimulates neuroendocrine hyperplasia that is specific to the stomach. However unlike the islets and pituitary, glial-directed *Men1* deletion was insufficient to convert the hyperplastic endocrine cells into NETs in this model.

**Figure 3.**
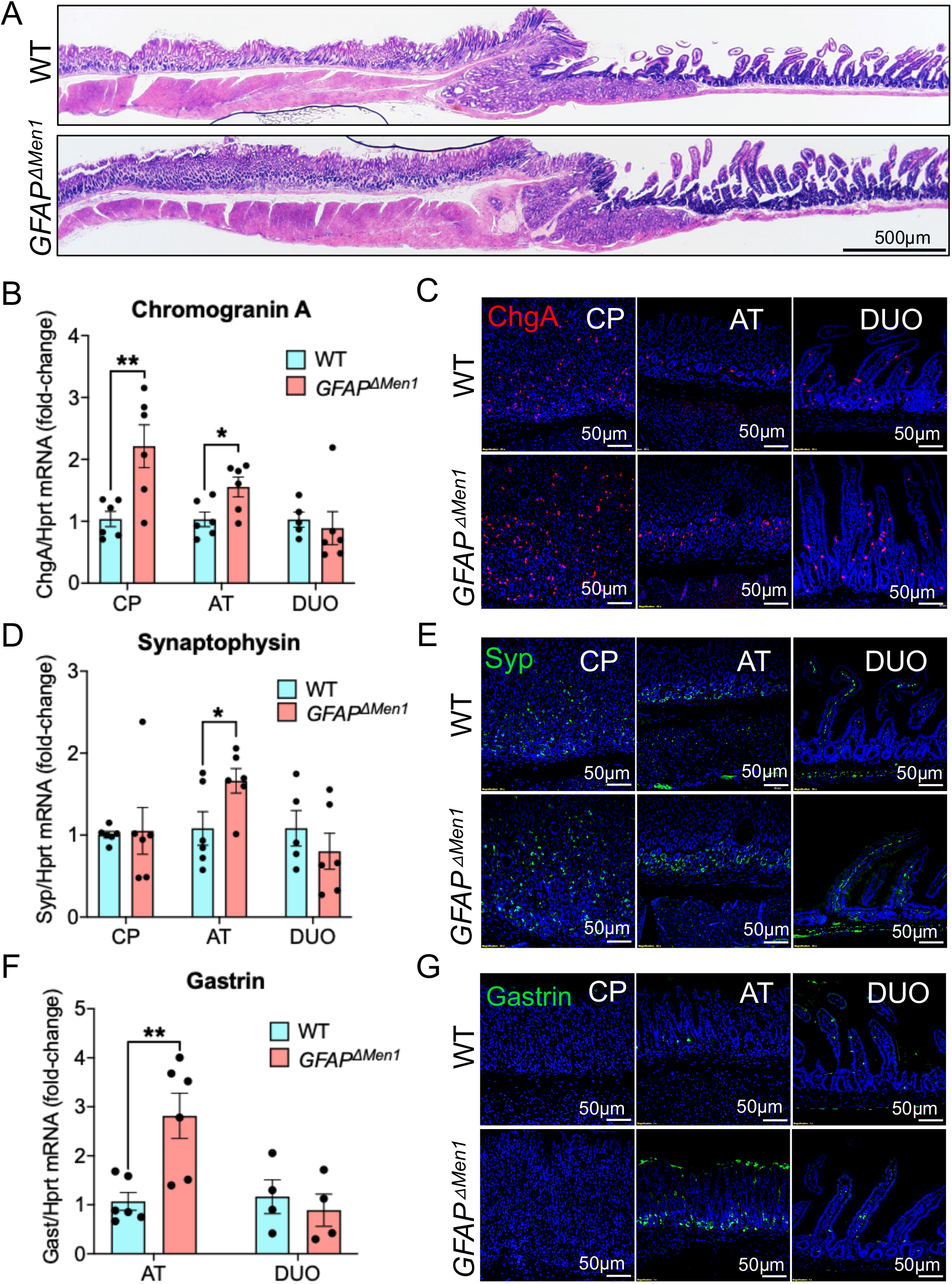
GFAP-directed deletion of Men1 stimulates hyperplastic reprogramming of the gastro-intestinal epithelium and promotes neuroendocrine cell hyperplasia. (A) Representative low power H&E stained images of WT and *GFAP*^*ΔMen1*^ antrum and the proximal duodenal mucosa at 8-9 months of age. (B) Expression of chromogranin A mRNA in WT and *GFAP*^*ΔMen1*^ corpus (CP), gastric antrum (AT) and duodenal mucosa (DUO). (C) Immunofluorescent staining of the corpus, antrum and duodenum for chromogranin A. (D) Synaptophysin mRNA expression in WT and *GFAP*^*ΔMen1*^ corpus, gastric antrum and duodenal mucosa. (E) Immunofluorescent staining of the corpus, antrum and duodenum for synaptophysin. (F) Gastrin mRNA and (G) Immunofluorescent staining for gastrin in the corpus, antrum, and duodenum. n=6 mice per group; * *p* < 0.05, ** *p* < 0.01 by unpaired Student’s T-test. Data are represented as mean ± SEM.

### Glial cell reprogramming occurs in *GFAP*^*ΔMen1*^ mice

Since NET development in the pituitary and pancreas coincided with an apparent loss of GFAP expression, we examined the impact of *Men1* deletion on glial cell fate *in vivo*. We generated *GFAP*^*ΔMen1*^ mice expressing a *GFAP Cre* tdTomato reporter (Figure 4A). Fluorescent imaging of both whole tissue mounts and frozen sections of the GI tract confirmed expression in enteric glial cells. Surprisingly, *ex vivo* fluorescent imaging of tissues from *GFAP*^*ΔMen1*^ mice revealed a ∼4-fold reduction in the tdTomato signal across the upper GI tract compared to wild type mice (Figure 4B and C). Fluorescent imaging indicated appropriate localization of the tdTomato signal to the lamina propria, submucosa and myenteric plexus ganglia (Figure 4D and E). Further imaging of frozen tissue sections confirmed near-loss of tdTomato fluorescence in the mucosa and lamina propria, while some signal was still retained in the submucosal and myenteric plexi (Figure 4E). Therefore, deletion of *Men1* in GFAP+ cells extinguished GFAP expression.

**Figure 4.**
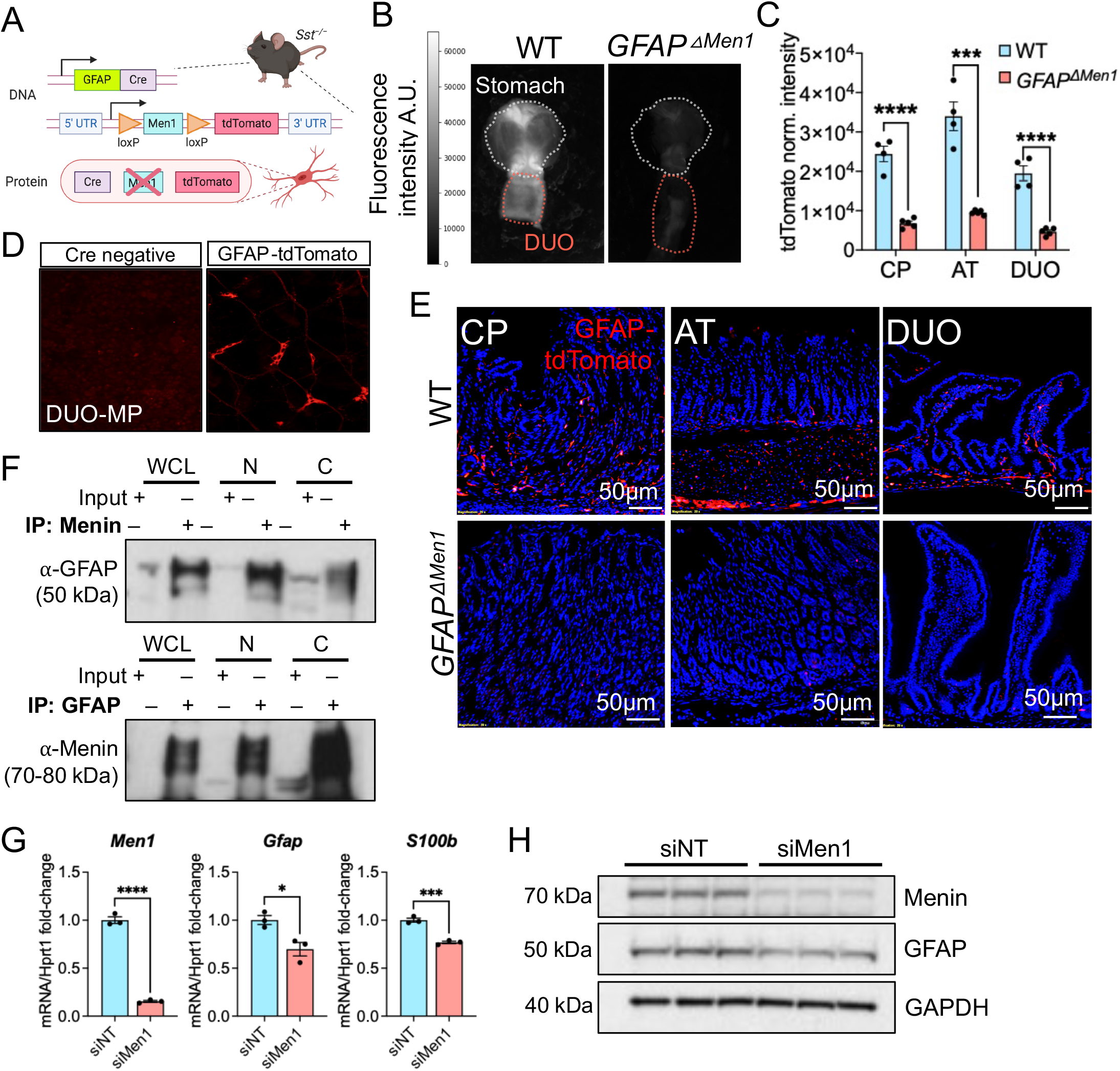
Conditional deletion of menin in GFAP^+^ cells stimulates loss of a mature glial cell phenotype. (A) Schematic of *GFAP*^*△Men1*^-tdTomato construct. (B) Widefield images and (C) quantitation of tdTomato signal in the stomach and duodenum of wild type and *GFAP*^*△Men1*^ mice expressing tdTomato reporter. n=4–5 mice per group; *** *p* < 0.001, **** *p* < 0.0001 by unpaired Student’s T-test. (D) Whole tissue mounts of proximal duodenum from *Cre* negative and tdTomato-expressing mice showing tdTomato fluorescence localized to the myenteric plexus (MP). (E) Representative images of cryosections of corpus (CP), gastric antrum (AT), and proximal duodenum (DUO) from wild type and *GFAP*^*△Men1*^ mice expressing tdTomato. (G) Co-immunoprecipitation (Co-IP) of menin and GFAP from rat EGC whole cell lysate (WCL), and nuclear (N) and cytoplasmic (C) fractions followed by western blot for GFAP and menin, respectively. Input is 5–10% of lysate used for IP. (H) Expression of *Men1* and the glial transcripts *Gfap* and *S100b* following siRNA-mediated *Men1* silencing in cultured rat enteric glial cells. n=3 independent experiments; * *p* < 0.05, *** *p* < 0.001, **** *p* < 0.0001 by unpaired Student’s t-test. Data are represented as mean ± SEM. (I) Western blot of menin and GFAP in whole cell lysates following 72h *Men1* silencing in rat enteric glial cells.

A prior study showed that GFAP and menin proteins interact in astrocytes ^[23]^. To test whether this interaction also occurs in glial cells we performed co-immunoprecipitation using extracts from a rat enteric glial cell line (EGC) and confirmed interactions between GFAP and menin in whole cell lysates as well as nuclear and cytoplasmic fractions (Figure 4F). Although intermediate filaments including GFAP are elaborate cytoskeletal structures residing in the cytoplasm, their expression also occurs in the nucleus and peri-nuclear regions of various cell types ^[25–27]^. Similarly, the subcellular localization of menin is dynamic with its movement in and out of the nucleus regulated by multiple factors ^[14,28,29]^. Accordingly, menin and GFAP interactions occurred in both the nuclear and cytoplasmic cellular compartments (Figure 4F). Moreover siRNA-mediated *Men1* silencing, also decreased S100b, another mature glial cell marker along with GFAP (Figure 4G and H), indicating a role for menin in regulating glial cell identity by modulating GFAP.

### Transcriptome-wide analysis of gastric neuroendocrine hyperplasia and NENs supports glial-to-neuroendocrine reprogramming

Since glial-specific deletion of menin was sufficient to drive NEN formation in the pituitary and pancreas but not in the stomach or intestine, we used RNA-Seq to identify the molecular pathways that might contribute to the transition from neuroendocrine hyperplasia to NEN development. *GFAP*^*ΔMen1*^ pitNETs, gastric antrum, and their respective wild type (WT) counterparts were submitted for bulk RNA sequencing. Principal Component Analysis confirmed a higher degree of divergence between pitNETs and normal WT pituitaries compared to hyperplastic *GFAP*^*ΔMen1*^ and WT antra (Supplementary Figure 1A). Accordingly, the number of significant differentially expressed genes (DEGs) was ten-times higher in pitNETs compared to antral tissues (Figure 5A and B). As anticipated, a number of genes related to neuroendocrine differentiation were upregulated in both datasets, e.g. *Grp, Vip, Syp, and Ncam1/2* in antra and *Prl, Chgb, Eno2, Cckbr, Vip*, and *Ascl2* in pitNETs (Figure 5C and D). Genes associated with the neuroglial lineage were significantly upregulated in *GFAP*^*ΔMen1*^ tissues compare to WT controls (Figure 5E and F). Furthermore, a majority of upregulated antral genes were correlated with neural differentiation and acquisition of a neuronal cell fate, including *Ngfr, Elav3/4, Nrsn1/2, Phox2b, Nos1, Tubb3, Uchl1, Ret*, and *Hoxb* (Figure 5E). PitNETs also exhibited increased expression of neural-specific transcripts, e.g. *Elav3/4, Nrsn1/2*, and *Tubb3* and this coincided with significant downregulation of astroglial lineage genes, e.g. *Gfap, Slc1a3, Cd40*, Aldh1l1, *Aqp4, Olig2, P2ry12, Sox9*, and *Notch1*, (Figure 5F). Furthermore, pitNETs exhibited increased expression of neural crest cell transcripts related to stemness, including *Vim, Fabp7*, and *Sox11*, as well as genes involved in cell cycle inhibition and differentiation (e.g. *Cdkn1a, Cdkn2a*, and *Cend1*), as well as tissue development (e.g. *Shh, Nog, Bmp2/7, Bdnf*, and *Ezh2*) (Figure 5F and Supplementary Figure 1B and C). Increased expression of genes related to NET differentiation and cellular senescence were validated by quantitative PCR of NETs and age-matched WT tissues (Supplementary Figure 1D and E).

**Figure 5.**
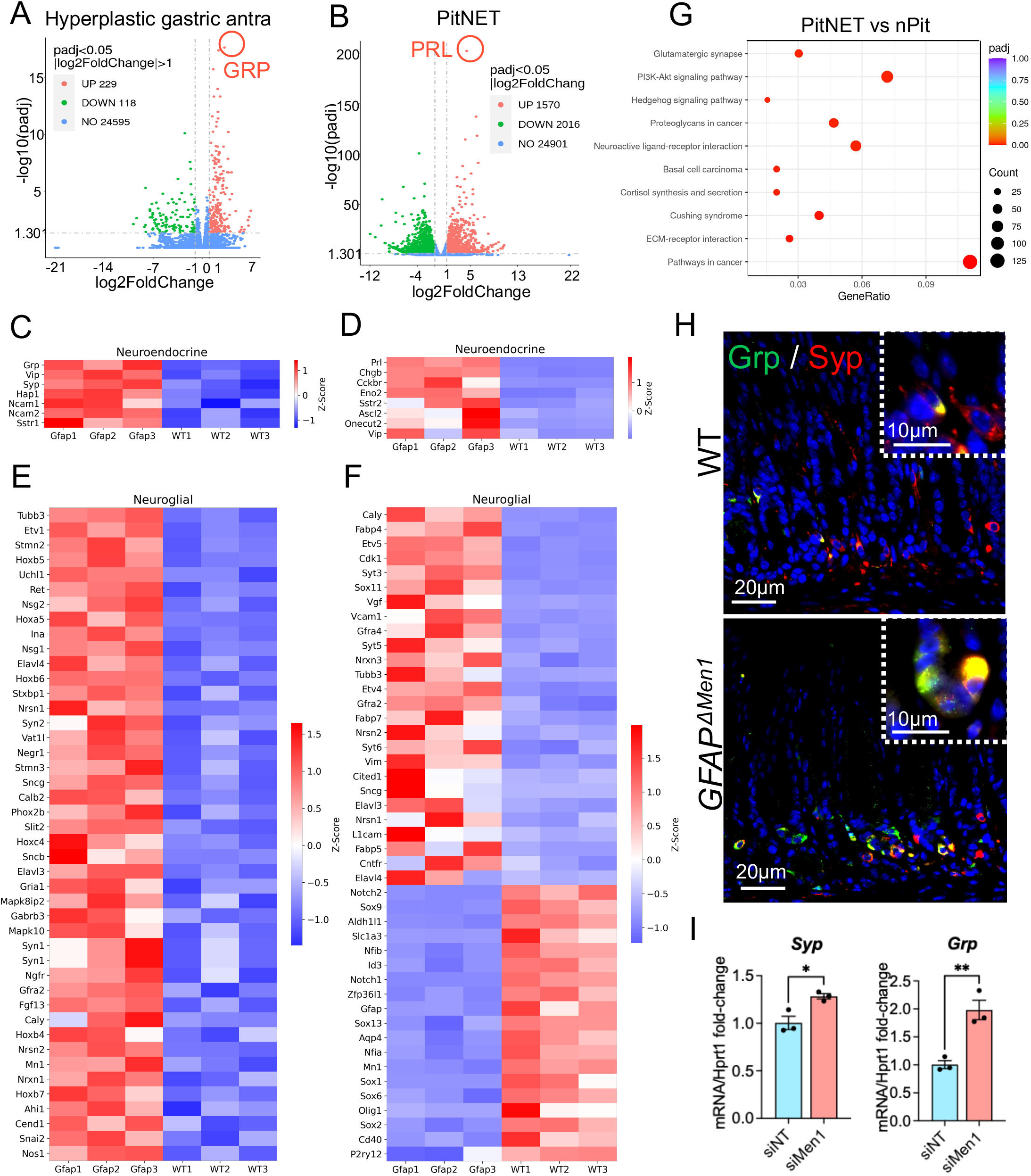
Transcriptome-wide analysis of gastric neuroendocrine hyperplasia and NENs identifies a role for glial-to-neural reprogramming in epithelial differentiation. (A) Volcano plot of significant DEGs in gastric antra of wild type and *GFAP*^*△Men1*^ mice. (B) Volcano plot of significant DEGs in pooled wild type pituitaries and pitNETs of *GFAP*^*△Men1*^ mice. (C) Heatmaps of significant DEGs mapped to neuroendocrine differentiation in *GFAP*^*△Men1*^ gastric antra and (D) pitNETs. (E) Heatmap of significant DEGs mapped to the neuroglial lineage in *GFAP*^*△Men1*^ gastric antra and (F) pitNETs compared to litter-mate wild type controls. (G) KEGG Ontology Pathway analysis of gastric antra from wild type and *GFAP*^*△Men1*^ mice showing the number of genes mapped to enriched pathways and their level of statistical significance. (H) Immunofluorescent staining for gastrin-releasing peptide (Grp) and synaptophysin (Syp) in WT and *GFAP*^*△Men1*^ gastric antra, shown as green and red respectively. Insets: colocalization of Grp and Syp in the gastric mucosa (yellow). (I) Quantitation of synaptophysin and Grp mRNA following 48h siRNA-mediated silencing of *Men1* in rat enteric glial cells. n=3 independent experiments; * *p* < 0.05 and ** *p* < 0.01 by unpaired Student’s t-test. Data are represented as mean ± SEM.

Both datasets showed an enrichment of genes involved in neuroactive ligand receptor interaction and synaptic signaling pathways (Figure 5G and Supplementary Figure 1F). A number of genes upregulated in pitNETs were mapped to the Hedgehog signaling pathway, which was consistent with increased expression of Shh mRNA and ciliary motor proteins known to facilitate transduction of Hedgehog signaling (Figure 5G and Supplementary Figure 1D and G). Further, increased expression of neuroendocrine and neural-lineage genes in *GFAP*^*ΔMen1*^ antral extracts coincided with downregulation of cytokeratins and immune-related transcripts (Supplementary Figure 1H and I).

Enteric neurons express gastrin-releasing peptide (Grp) ^[30]^, which is the most significantly upregulated transcript in *GFAP*^*ΔMen1*^ gastric antra, providing a direct mechanism for increased antral gastrin levels in these mice. To confirm that GFAP-directed deletion of *Men1* stimulates Grp expression leading to G cell expansion, we stained the antra of *GFAP*^*ΔMen1*^ mice for Grp. As anticipated, *GFAP*^*ΔMen1*^ mice exhibited increased antral expression of Grp. Furthermore, Grp expression co-localized with Syp in the epithelial mucosa (Figure 5H). To assess whether loss of menin in enteric glial cells (EGCs) directly stimulates the expression of Grp and Syp, *Men1* was knocked down in rat EGCs, which induced both Grp and Syp mRNA expression (Figure 5I). Taken together, these results demonstrate that *Men1* deletion suppresses the mature glial phenotype in favor of a neuroendocrine phenotype, perhaps through loss of GFAP.

### Impairment of Sonic hedgehog (Shh) signaling in enteric glial cells attenuated neuroendocrine cell hyperplasia in *GFAP*^*ΔMen1*^ mice

Hedgehog signaling mediated by primary cilia is known to promote premature differentiation of neural progenitor cells toward the neuronal lineage ^[31]^. As *GFAP*^*ΔMen1*^ pitNETs exhibited significantly elevated Shh transcript levels, we investigated whether loss of *Kif3a*, a ciliary motor protein required for Shh signal transduction, blocks *GFAP*^*ΔMen1*^ glia from developing a neuroendocrine phenotype. To further stimulate gastric neuroendocrine hyperplasia, *GFAP*^*ΔMen1*^ mice were crossed onto a somatostatin null (*Sst*^*–/–*^) background. Sst suppresses hormone secretion, including gastrin, thus removal of *Sst*-mediated feedback inhibition increases gastrin levels ^[14]^. The effect of *Kif3a* deletion on neuroendocrine hyperplasia was accomplished by breeding *GFAP*^*ΔMen1*^*;Sst*^*–/–*^ mice onto a *Kif3a*^*FL/FL*^ genetic background (*GFAP*^*ΔMen1;ΔKif3a*^) (Figure 6A). Shortened primary cilia were shown by immunostaining for acetylated tubulin-positive hair-like projections on enteric glia. Compared to WT mice, *GFAP*^*ΔMen1;ΔKif3a*^ mice exhibited shortened primary cilia on GFAP^+^ glial cells in the gastric antrum (Figure 6B). Hedgehog signaling is mediated through activation of the Gli1/2/3 family of transcriptional effectors ^[32]^. Thus, we assessed whether deleting *Kif3a* affects the expression of Gli2, a known suppressor of gastrin gene expression in antral G cells ^[33]^. Whereas GFAP-directed *Men1* deletion increased antral Gli2 expression, loss of *Kif3a* reduced the expression of Gli2 protein and transcript (Figure 6C and D). We next assessed whether *GFAP*^*ΔMen1;ΔKif3a*^ mice showed reduced GFAP-tdTomato fluorescence compared to *GFAP*^*ΔMen1*^ mice with intact primary cilia. As expected, loss of GFAP-tdTomato fluorescence was reversed in the *GFAP*^*ΔMen1;ΔKif3a*^ mice (Figure 6E and F). *Kif3a* deletion was associated with a trending increase in antral *Gfap* mRNA expression (Figure 6G). To determine the effect of *Kif3a* deletion on enteroendocrine cell composition, we analyzed the neuroendocrine -related transcripts in the antrum and proximal duodenum after sequential deletion of *Sst, Men1*, and *Kif3a* genes. Compared to wild type and *Sst* null mice, deleting *Men1* significantly increased the expression of ChgA and Syp mRNA in the gastric antra, as anticipated (Figure 6H and I). Deleting *Kif3a* in the presence of *Sst* and *Men1* deletions inhibited the previously observed increase in antral neuroendocrine transcripts induced antral expression of gastrin transcript and gastrin peptide in *GFAP*^*ΔMen1*^ mice. The induction was mitigated after *Kif3a* was deleted in (Figure 6J and K). Thus, blockade of Shh-mediated signaling in enteric glia attenuated neuroendocrine cell hyperplasia in the distal stomach.

**Figure 6.**
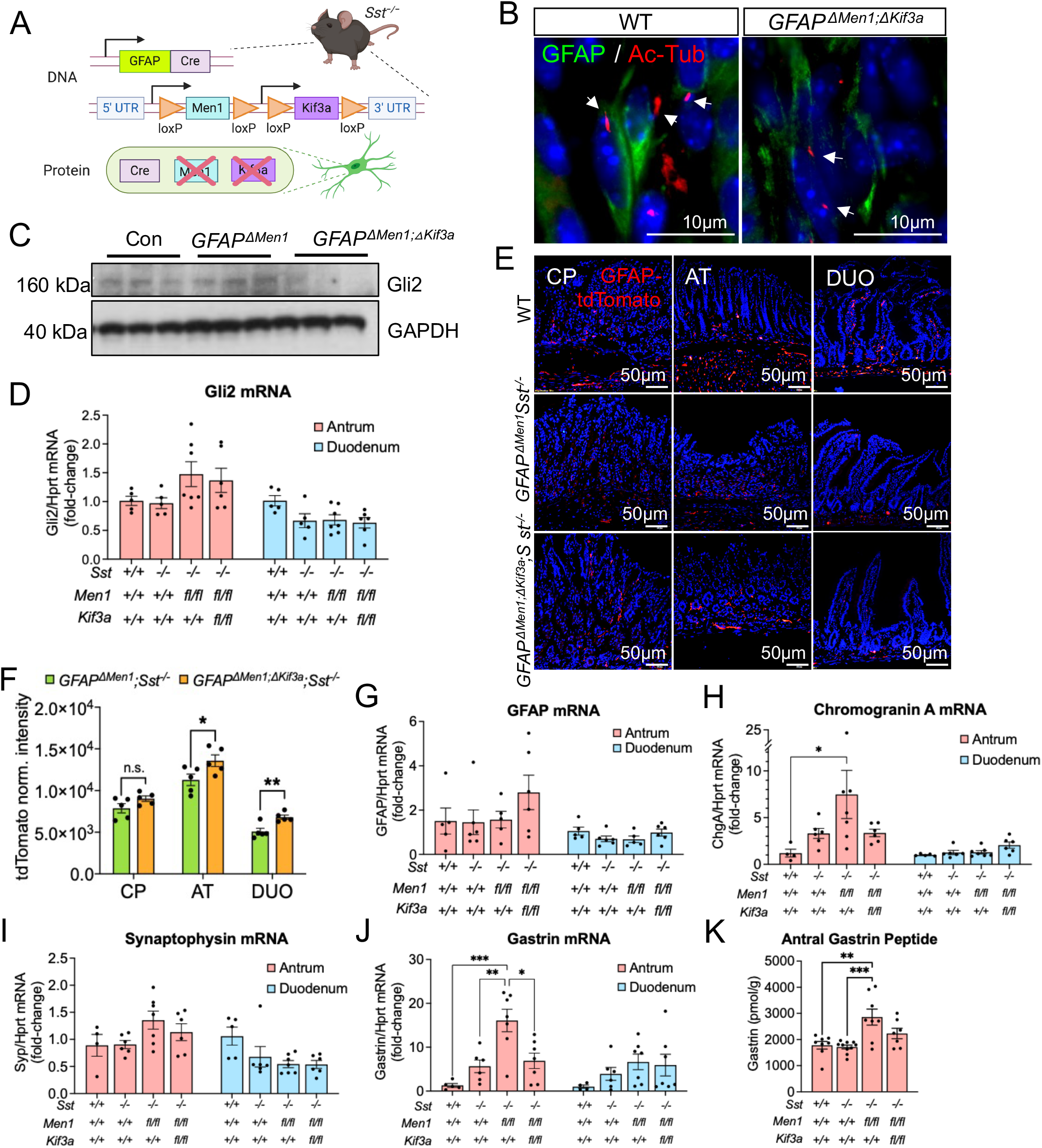
Impairment of Sonic hedgehog (Shh) signaling in enteric glial cells attenuates neuroendocrine differentiation in *GFAP*^*△Men1*^ mice. (A) Schematic of *GFAP*^*ΔMen1;ΔKif3a*^*;Sst*^*-/-*^ mouse derivation. (B) Immunofluorescent staining for acetylated-tubulin and GFAP in the gastric antra of wild type and *GFAP*^*ΔMen1;ΔKif3a*^*;Sst*^*-/-*^ mice. Arrows indicate primary cilia marked by acetylated tubulin. (C) Western blot of Gli2 protein in gastric antral lysates of *Sst*^*-/-*^ (Con), *GFAP*^*△Men1*^*;Sst*^*-/-*^, and *GFAP*^*ΔMen1;ΔKif3a*^*;Sst*^*-/-*^ mice. n=3 mice. (D) Gli2 mRNA in gastric antra and duodenal mucosa of respective genotypes. n=4–7 mice per group. (E) Representative images of cryosections of corpus (CP), antrum (AT), and proximal duodenum (DUO) from wild type, *GFAP*^*△Men1*^*;Sst*^*-/-*^ and *GFAP*^*ΔMen1;ΔKif3a*^*;Sst*^*-/-*^ mice expressing tdTomato. (F) Quantitation of relative tdTomato fluorescence intensity following whole tissue ex vivo imaging. * *p* < 0.05, ** *p* < 0.01 by unpaired Student’s T-test. (G) Quantitation of GFAP, (H) Chromogranin A, (I) Synaptophysin, and (J) Gastrin mRNA in the gastric antrum and duodenum of different groups. n=4–7 mice per group; * *p* < 0.05, ** *p* < 0.01, *** *p* < 0.001 by unpaired Student’s T-test. (K) Expression of antral gastrin peptide amongst the different genotypes. n=7–8 mice per group; ** *p* < 0.01, *** *p* < 0.001. All data are represented as mean ± SEM.

## DISCUSSION

Progress in understanding GEP-NET pathogenesis is impeded by a lack of comprehensive *in vivo* models, in part due to tissue heterogeneity from which the neoplasms arise, and the absence of consistent driver mutations preceding malignancy ^[34]^. We report here that conditional deletion of the tumor suppressor protein *Men1* (menin) in GFAP^+^ glial cells induced neuroendocrine cell hyperplasia and the development of neuroendocrine tumors in the pituitary and pancreas.

Neuroendocrine cell hyperplasia and NET development coincided with an unexpected loss of the mature glial cell phenotype and induction of neuronal and neuroendocrine-lineage genes in stomach, pancreas and pituitary. We demonstrated that deletion of *Men1* induced loss of GFAP in the GI tract and in NETs by immunostaining, qPCR, and through tracking of GFAP-tdTomato positive cells. In contrast to NETs which exhibited a global reduction in GFAP staining, tdTomato signal, and mRNA expression, the gastric antra of *GFAP*^*ΔMen1*^mice showed reduced GFAP-reporter expression but did not exhibit significant loss of GFAP at the transcript level. This observation may be explained in part by the fact that menin interacts and colocalizes with GFAP ^[23]^. Hence, it remains a possibility that conditional deletion of menin in GFAP^+^ glial cells disrupts proper localization of GFAP leading to altered glial cell identity creating a microenvironment sufficient for tumor formation in the islet pancreas and pituitary but not in the gastric antrum or duodenum.

Similar to previous reports of *Men1*-deleted mouse models, we did not observe the development of small intestinal gastrinomas, suggesting that additional genetic or microenvironmental factors are required to stimulate neoplastic transformation in these tissues ^[12,35–39]^. To identify molecular features that might inform the transition from neuroendocrine hyperplasia to NET development, we compared the transcriptomes of hyperplastic antral tissues with well differentiated NETs arising in this model. While glial-directed *Men1* deletion induced neural differentiation pathways at both tissue sites, NETs exhibited loss of more astroglial-lineage genes, e.g. *Gfap, Aldh1l1, Slc1a3, Aqp4, and Olig1* and downregulation of factors involved in directing astrogliogenesis, e.g. *Nfia, Nfib, Zfp36l1, Id3*, and *Sox9* ^[40–44]^. Thus, the transition from a glial-to-neuronal cell phenotype appears to promote the hyperplastic neuroendocrine and tumor development. In support of this, NETs exhibit increased expression of genes associated with neural stem and progenitor cell status, e.g. *Fabp7, Vim, Sox11*, and *Hoxb* genes, and upregulation of neural crest-secreted factors that favor neurogenesis and restrict a glial cell fate, e.g. *Bmp2, Bmp7*, and *Shh* ^[45–51]^. A number of these signaling factors are components of the Hedgehog signaling pathway, known to play a role in directing neuronal differentiation at the expense of restraining gliogenesis and glial cell maturation ^[31]^. A role for menin in this process is highlighted by the fact that menin is a known epigenetic repressor of both canonical and non-canonical Hedgehog signaling ^[52,53]^. Moreover, these studies demonstrate that blockade of Hedgehog signaling attenuate islet cell proliferation in a *Men1*-mediated insulinoma mouse model ^[53]^. Consistent with these reports, we found that attenuation of Shh signaling by disrupting primary cilia on glial cells reduced neuroendocrine hyperplasia by restricting neural differentiation of *Men1*^*FL/FL*^ glial cells.

Enteric glia exhibit a high degree of cellular plasticity within the context of their specific microenvironment ^[38, 54]^. Indeed, they have been identified as a source of neural progenitor cells and are able to de-differentiate and trans-differentiate to other cell lineages under explicit physiological and *in vitro* conditions ^[15,16,55–58]^. However, it remains unknown whether neural crest cells or their progeny can undergo context-specific reprogramming, e.g., by acquiring *MEN1* mutations, and giving rise to neuroendocrine cells with hormone-secreting capabilities. Evidence to support glial or astrocytic reprogramming would challenge the long-standing paradigm that neuroendocrine tumors in the GI tract develop from enteroendocrine cells that originate from Lgr5^+^ stem cells of endodermal origin ^[59]^. Previous work by our group demonstrated that non-cell autonomous loss of menin protein in enteric glial cells induces gastrin hormone expression ^[13,14]^. Furthermore, human duodenal NETs are known to express glial cell markers, with some cells within the tumor exhibiting expression of both the neuroendocrine marker Syp and the glial-specific protein S100b ^[14,60]^. We recently used digital spatial profiling to characterize neuroglial features in a small subset of human duodenal NETs ^[60]^. Tumors exhibited reduced expression of mature neuronal and glial cell markers compared to the adjacent Brunner’s glands from which 60% of these tumors are reported to originate ^[9]^. Collectively, these observations raise the potential for the presence of a potential hybrid transition state, in which reprogrammed enteric neural crest-derived cells escape a mature neuroglial lineage and acquire a neuroendocrine phenotype.

In identifying molecular features unique to the tumor-naïve hyperplastic antrum, we focused on a subset of genes enriched in our study and previously reported to be upregulated in human duodenal gastrinomas (DGASTs), including *Hap1, Mn1*, and *Uchl1* ^[60]^. Among these, the neuronal protein Huntingtin-associated protein (Hap1) is also expressed by enteroendocrine cells (EECs) within the GI tract and was recently reported to be expressed in G cells but not in other EEC lineages of the pyloric antrum ^[61]^. This is consistent with our observation of increased *Hap1* gene expression and higher antral gastrin and Grp expression levels. In addition, *GFAP*^*ΔMen1*^ mice exhibited increased expression of Meningioma 1 (Mn1), a non-glial neural progenitor marker that is strongly associated with primitive neural ectodermal tumors of neural crest cell origin ^[62]^. Similarly, Ubiquitin C-Terminal Hydrolase L1 (Uchl1) is a pan-neuronal marker that is also expressed by neural progenitor cells of the enteric nervous system and by neuroendocrine cells ^[63–65]^.

Taken together, enrichment of these neural lineage-specific genes in both the hyperplastic antra of *GFAP*^*ΔMen1*^ mice and human DGASTs arising from the Brunner’s glands suggests concordance between the human Brunner’s glands and the pyloric antrum. A recent study showed that transplanted human antral organoids co-cultured with human ENCCs induced the formation of highly differentiated epithelium analogous to the Brunner’s glands ^[66]^. Subsequent analysis of ENCCs identified elevated expression of the Shh-induced posteriorizing factors Bmp4 and Bmp7, and inhibition of these factors in ENCCs attenuated Brunner’s gland formation in organoid co-cultures ^[61]^. Consistent with our observations, ENCCs and their progeny carry the potential to direct epithelial differentiation in favor of an endocrine phenotype, and these events are strongly influenced by Shh-mediated signaling. Importantly, our observations prompt further investigation into both neuroglial cell autonomous and non-cell autonomous mechanisms presaging these outcomes.

In summary, we report the development of a glial-directed mouse model of human MEN1 syndrome with the aim of defining the contribution of neural crest cell reprogramming in neuroendocrine cell differentiation and tumorigenesis. By addressing a previously uncharacterized compartment of GEP-NETs, this study carries the potential to identify a unique cell-of-origin for these neoplasms. The importance of these discoveries is emphasized by the fact that GEP-NETs comprise remarkably diverse neoplasms that vary in location, mutational profile, and response to therapy. Such heterogeneity may in part be explained by divergent cellular origins for NETs as a function of different tissue sites. Previous work centered on mapping their unique transcriptional signatures suggests the potential for neuroglial reprogramming in the development of human DGASTs. Defining the cell-of-origin and the events preceding neoplastic transformation will be critical to informing molecular signaling pathways that can then be targeted therapeutically.

## MATERIALS AND METHODS

All authors had access to the study data and had reviewed and approved the final manuscript.

### Animal Studies

All animal studies received approval by the University of Arizona Institutional Animal Care and Use Committee and conform to the Animal Research: Reporting of In Vivo Experiments guidelines. *GFAP-Cre* transgenic mice on a C57BL/6J genetic background were purchased from Jackson Laboratories and bred onto a *Men1*^*FL/FL*^ background (*GFAP*^*ΔMen1*^) to conditionally delete *Men1* in GFAP-expressing cells. A subset of wild type and *GFAP*^*ΔMen1*^ mice were bred onto a somatostatin-null background (*Sst*^*-/-*^). *GFAP*^*ΔMen1*^*;Sst*^*-/-*^ mice were further bred onto a *Kif3a*^*FL/FL*^ background to conditionally delete *Kif3a* under the control of the *Gfap* promoter (*GFAP*^*ΔMen1;ΔKif3a*^*;Sst*^*-/-*^). Wild type, *GFAP*^*ΔMen1*^, *GFAP*^*ΔMen1*^*;Sst*^*-/-*^, and *GFAP*^*ΔMen1;ΔKif3a*^*;Sst*^*-/-*^ mice were also bred to mice carrying a *lox-Stop-lox-tdTomato* sequence to selectively express the tdTomato fluorescent reporter in GFAP^+^ cells. Mice were necropsied at 8–9 months of age and between 13–24 months of age. All histological characterization and downstream comparisons were made using littermate controls.

### Cell and Tissue Culture

Primary tumor neurospheres were generated from an 18 mo-old female *GFAP*^*ΔMen1*^ mouse presenting with a large pituitary prolactinoma. A four-by-two millimeter piece of tissue was cut from the tumor and minced into 1 mm pieces on ice. The tissue was transferred to a 10 mL solution of Dispase/DNase solution warmed to 37°C (0.1% dispase, 0.01% DNase, 10 mM HEPES, DPBS without calcium and magnesium) and incubated in a shaking water bath (60 rpm) at 37°C for 80 minutes. Half-way through the incubation, the tissue was shaken manually three times to facilitate dissociation. The tissue was triturated 10 times with a 10 mL serological pipette, then filtered through a 40 μm cell strainer before centrifuging at 123 x g for 10 min at 4°C. The cell pellet was resuspended in Neurosphere Media consisting of DMEM/F12 (ThermoFisher, Waltham, MA), 1X GlutaMAX (Invitrogen, Carlsbad, CA), 1X N2 (Gibco, Waltham, MA), 1X B-27 (Gibco), 100U Penicillin-streptomycin (Invitrogen), 10 mM HEPES (Invitrogen), 30 ng/mL mouse recombinant FGF-10 (Biolegend, San Diego, CA), and 30 ng/mL recombinant human EGF (R&D Systems, Minneapolis, MN). Cells were plated in a non-tissue culture treated 6-well plate and incubated overnight at 37°C. Media was changed every 2-3 days by collecting neurospheres, centrifuging, and reseeding the pellet in fresh medium.

A rat enteric glial cell line was purchased from ATCC (#CRL-2690) and grown in DMEM supplemented with 10% Fetal Bovine Serum (FBS) and 100U Penicillin-streptomycin (EGC/PK060399egfr, ATCC, Manassas, VA). Upon reaching 30-50% confluency, glial cells were switched to 5% FBS-containing media and used for small interfering RNA (siRNA) experiments. Both the non-targeting siRNA and Men1 siRNA consisted of a SMARTpool of four ON-TARGETplus siRNA constructs (#L-090784-02-0005 for *Men1* pool and #D-001810-10-05 for Non-targeting pool, Dharmacon Horizon Discovery, Lafayette, CO). Cells were transfected with 25 nM siRNA using Lipofectamine 3000 without p3000 reagent according to manufacturer instructions (ThermoFisher). Following 48 to 72h post-transfection, cells were harvested for downstream RNA or protein extraction.

### Immunohistochemistry and Immunofluorescent Staining

Mouse tissues were fixed overnight at 4°C in 4% paraformaldehyde. Tissues were paraffin-embedded, cut into 5 μm sections and placed on frosted glass slides. Slides were deparaffinized in three xylene washes, then rehydrated in 100%, 90%, and 70% ethanol washes. Slides were washed in TBS with 0.05% Tween-20 (TBST) before proceeding with antigen retrieval. Slides were boiled in 1X sodium citrate buffer pH 6.0 for 30 min and allowed to cool to room temperature (RT) for 20 min prior to washing with TBST. Slides were incubated for 1h at RT in blocking buffer consisting of 10% donkey serum, 1% Bovine Serum Albumin (BSA), 0.1% Triton X-100 in TBST. In some instances, tissues were permeabilized with 0.5% Triton X-100 for 5 min at RT prior to blocking. Tissues were incubated in primary antibody overnight at 4°C. Primary antibodies were prepared in blocking solution according to the dilutions listed in STAR Methods Table 1. Following overnight incubation, slides were washes in TBST and incubated in Alexa Fluor-conjugated secondary antibodies (Molecular Probes, Eugene, OR; Invitrogen) diluted 1:500 in 1% BSA and TBST for 1h at RT in the dark. Slides were washed in TBS before mounting with #1.5 coverslips using Prolong Gold anti-fade mounting medium with DAPI (Life Technologies, Rockville, MD). Immunofluorescence was visualized using the Olympus BX53F epifluorescence microscope (Center Valley, PA).

For immunofluorescent staining of tumor neurospheres, neurospheres were collected and centrifuged. The pellet was resuspended in 3.7% paraformaldehyde and transferred to a 24-well 0.17mm glass bottom plate. Neurospheres were fixed for 15 min at RT, then washed with 1X TBST. Neurospheres were permeabilized with 0.5% triton X-100 for 5 min at RT, washed, then blocked in 20% donkey serum, 1% BSA and 0.1% triton in TBST for 30 min at RT. Primary antibodies were diluted in 5% donkey serum 1% BSA, 0.1% triton in TBST and added to wells for overnight incubation at 4°C. Primary antibodies and dilutions are listed in Methods Table 1. Neurospheres were washed thoroughly with TBST and incubated for 1h at RT in Alexa Fluor-conjugated secondary antibodies diluted 1:500 in 1% BSA and TBST. Neurospheres were washed with TBS, then stored in TBS containing Prolong gold antifade mounting medium with DAPI. Samples were imaged using a Nikon Ti2-E inverted microscope with a Crest X-Light V2 L-FOV Spinning Disk Confocal and a Photometrics Prime 95B-25MM Back-illuminated sCMOS Camera (Nikon, Tokyo, Japan).

Mouse tissues with endogenous tdTomato reporter expression were fixed in 4% paraformaldehyde for 1h on ice before dehydrating overnight at 4°C in a solution of 30% sucrose-PBS. Tissues were then embedded in Tissue-Tek Optimal Cutting Temperature (OCT) compound (Sakura Finetek, Torrance, CA) and frozen on dry ice. OCT blocks were cryosectioned to a 10 μm thickness and melted onto frosted glass slides prior to mounting with Prolong Gold anti-fade mounting medium containing DAPI. TdTomato signal was visualized using the Olympus BX53F epifluorescence microscope. Images drawing comparisons between groups were acquired with identical laser acquisition settings (Center Valley, PA).

### Quantitative PCR

RNA was extracted from frozen mouse tissues using the Purelink RNA Extraction Kit (Invitrogen). Tissues were homogenized with a rotor-stator homogenizer in RNA lysis buffer containing 1% β-mercaptoethanol and RNA was isolated following manufacturer’s instructions. Extracted RNA was treated with ezDNase to remove any residual genomic DNA and then prepared for cDNA synthesis using Superscript VILO IV Master Mix according to manufacturer’s instructions (Invitrogen). Quantitative real-time PCR was performed using PowerUp SYBR Green Master Mix (Invitrogen) with 20 ng cDNA added to each reaction. Predesigned forward and reverse primer sets were purchased from Integrated DNA Technologies (IDT, Coralville, IA) and used at a final 500 nM concentration. Quantitative PCR was performed using the QuantStudio 3 Real-Time PCR System (Applied Biosystems, Waltham, MA) with the following cycling conditions: 2 min at 50°C, 2 min at 95°C, denaturing step for 1 sec at 95°C, extension and annealing for 1 min at 60°C, followed by a dissociation melt curve stage to confirm primer specificity. All forward and reverse primers were purchased as validated predesigned PrimeTime qPCR Assay primer sets used for SYBR® Green dye (Integrated DNA Technologies, Coralville, IA).

### *Ex vivo* Fluorescence Imaging

At 8–9 mos of age, *GFAP Cre-tdTomato* mice were euthanized following overnight fasting and the stomach and proximal duodenum were collected for *ex vivo* imaging prior to fixation for OCT embedding. The stomach and proximal duodenum were opened along the greater curvature and contents were flushed with cold PBS. Both tissues were placed in cold PBS with mucosa facing up and imaged in a dark room using a wide field fluorescence imaging system. Illumination was provided by a 300W xenon arc lamp source (LS-OF30, Sutter instruments, Novato CA). Illumination light was filtered with a 554 nm excitation filter with 23 nm bandwidth (FF01-554/23-25, Semrock Inc., Rochester NY) and delivered using a fiber bundle (LLG, Sutter Instruments). Images were collected with a ½” CCD Monochrome Camera (EO-1312M, Edmund Optics, Barrington NJ) with a 35 mm fixed focal length lens (#59-872, Edmund Optics) and a 594 nm long-pass filter (BLP01-594R-25, Semrock) mounted on a rigid arm above the sample plane. Images were collected with an exposure time of 60 ms and saved in 16 bit .tif format. Each day, a flat-field image was collected by imaging a white diffuse reflectance target (#58-609, Edmund Optics) to correct for spatial non-uniformity using unfiltered illumination. Additionally, a power measurement was collected for the filtered illumination light at the sample plane using a power meter (S120C, Thorlabs, Newton NJ) to adjust for day-to-day variations of light source intensity. The collected images were first normalized through division by the flat-field image and the light source power measured on the day of sample acquisition. Regions of interest were then manually drawn around the forestomach, corpus, antrum and duodenum using ImageJ. Fluorescence intensity was calculated in the region of interest and averaged over the number of valid pixels. Any pixels with 5% of saturation (e.g., digital value of 65535) were discarded from the analysis.

### RNA Sequencing

RNA was extracted from three *GFAP*^*ΔMen1*^ PNETs, pitNETs, and the pyloric antrum using the Purelink RNA Extraction kit as previously described (Invitrogen). Pancreas, pituitary, and antra of age-matched wild type mice were used as controls. Due to its small size, three normal pituitaries were pooled for each sample, and a total of three samples were submitted for sequencing (i.e. nine wild type pituitaries total). RNA was processed on the bioanalyzer for quality control prior to proceeding with mRNA library construction and bulk RNA sequencing. Due to degradation of pancreas tissues by pancreatic enzymes, only antral extracts and pituitary samples were usable for downstream library construction and sequencing. Samples were sequenced by Novogene using Ilumina sequencing platforms and the resulting calculated Fragments Per Kilobase of transcript sequence per Millions base pairs sequenced (FPKM) values were analyzed for differential gene expression, Principal Component Analysis, and Gene Ontology (GO) Enrichment Analysis. Differential expression between wild type and *GFAP*^*ΔMen1*^ groups was determined using the DESeq2 R package (1.20.0) using a negative binomial distribution model. *P* values were adjusted for multiple testing using the Benjamini and Hochberg’s method for controlling false discovery rate and genes with adjusted *p* value < 0.05 were considered significantly differentially expressed. The statistical enrichment of DEGs in KEGG pathways was tested using the clusterProfiler R package with correction for gene length bias. GO terms with a corrected *p* value < 0.05 were considered significantly enriched. Heatmaps of significant DEGs assigned to specific signatures of interest were generated using Python 3 using the open-source packages Seaborn and Pandas. For each gene, the Z-score was calculated and plotted as a function of color, with the color white being fixed to a Z-score of 0.

### Western Blot Analysis

Seventy-two hours following siRNA-mediated Men1 silencing, rat enteric glial cells were harvested by washing twice in cold DPBS and lysing in cold RIPA buffer containing phosphatase and protease inhibitors. Cells were collected by scraping, then homogenized by passing through a 20G needle syringe five times. The resulting lysate was centrifuged at 15,000 x g for 20 min at 4°C. The supernatant was collected and used for western blot analysis as follows. Lysates were prepared in reducing conditions in 1X SDS buffer with 5% β-mercaptoethanol and denatured by boiling at 95°C for 5 min. 15 μg of protein was loaded into a NuPage 4–12% Bis-Tris Mini Protein Gel and allowed to migrate for 1h at 140 V in 1X MOPS Gel Electrophoresis Buffer. Proteins were transferred onto a pre-wetted PVDF membrane using the iBlot2 transfer system (P0 setting, 7 min), then blocked in 5% BSA TBST for 1h at RT. Membranes were incubated overnight at 4°C in primary antibody diluted in blocking buffer at the dilutions listed in STAR Methods Table 1. Membranes were washed in TBST, then incubated for 1h at RT in a corresponding host IgG HRP-conjugated secondary antibody diluted 1:3000 in blocking buffer. Membranes were washed again in TBST and incubated in Pierce ECL reagent (ThermoFisher) for 1 min prior to developing on film in a dark room.

For analysis of tissue extracts, gastric antra of 8-9 mo-old mice were homogenized in ice-cold RIPA buffer using a rotor-stator homogenizer. Samples were centrifuged at 15,000 x g for 15 min and the supernatant (20μg) was evaluated for Gli2 protein expression (STAR Methods Table 1).

### Co-Immunoprecipitation Assay

Co-immunoprecipitation of menin and GFAP was performed in rat enteric glial cells. Prior to lysis, cells were collected in ice-cold DPBS, centrifuged, and washed once with cold DPBS. The cell pellet was resuspended in either RIPA buffer or cytoplasmic lysis buffer to generate a whole cell lysate and cytoplasmic extract, respectively. Buffers were supplemented with 1X HALT protease and phosphatase inhibitor. Whole cell lysates were prepared by passing through a 20G syringe 10X and centrifuging at 15,000 x g for 15 min at 4°C. Cytoplasmic extracts were pipetted up and down 20X, incubated on ice for 10 min, then vortexed for 15 sec prior to centrifuging at 15,000 x g for 5 min at 4°C. The supernatant was saved as the cytoplasmic fraction. The remaining pellet containing nuclear protein was washed once in cytoplasmic lysis buffer then resuspended in nuclear lysis buffer (1:4 v/v nuclear to cytoplasmic buffer ratio). The extract was incubated on ice for 20 min, vortexed vigorously, then centrifuged at 15,000 x g for 15 min at 4°C to remove chromatin. Protein concentration in freshly prepared lysates was evaluated by bicinchoninic assay. To form an antigen-antibody complex, 100-500 ug lysate was incubated with either Mouse anti-Menin monoclonal antibody (Bethyl Laboratories, #A500-003A, 1:100 v/v) or Rat anti-GFAP monoclonal antibody (Invitrogen, #13-0300, 1:100 v/v) overnight at 4°C. 50μL of prewashed Protein A/G agarose beads (Pierce, #20423) was added to each antigen-antibody complex and incubated overnight at 4°C with gentle agitation. Bead-complexes were pelleted at 3,000 x g for 2 min and washed three times with wash buffer. Bound proteins were eluted by mixing with 2X SDS loading buffer containing 5% % β-mercaptoethanol as a denaturing agent. Samples were incubated at 95°C for 10 min, vortexed, then centrifuged to remove beads. The remaining protein suspension was evaluated by western blot as previously described. Extracts precipitated using Mouse anti-Menin antibody were probed using Rabbit anti-GFAP antibody (Dako Agilent Technologies, #Z033429, 1:2000) whereas extracts precipitated with Rat anti-GFAP antibody were probed with Rabbit anti-Menin antibody (Bethyl Laboratories, #A300-105A 1:10000).

### Gastrin Enzyme Immunoassay

Gastrin peptide was extracted from the antra of 8–9 mo-old mice by boiling tissues in 200 μL of nuclease-free water and vigorously vortexing. The resulting solution was centrifuged to pellet debris and 50 μL of the supernatant was used for the Gastrin Enzyme Immunoassay according to manufacturer’s instructions (Sigma, #RAB0200).

### Statistical Analysis

Assays comparing two genotypes or groups were evaluated for statistical significance by applying unpaired Student’s t-test. All quantitative PCR data expressed as fold-change (2^-ddCt^) were log transformed to fit a normal distribution prior to performing statistical analysis. Comparisons consisting of three or more genotypes or groups were evaluated for significance using One-way Analysis of Variance (ANOVA), with Tukey post-test. Significance was determined as \**p* < 0.05, \*\**p* < 0.01, \*\*\**p* < 0.001, \*\*\*\**p* < 0.0001 using Graphpad Prism (v9) software.

**Methods Table 1.** List of primary antibodies and dilutions

| **Primary Antibodies** | **Vendor** | **Catalog #** | **Dilution/Application** |
| --- | --- | --- | --- |
| Menin (Rabbit pAb) | Bethyl Laboratories | A300-105A | 1:2,000 (IHC/IF with TSA)<br>1:10,000 (WB) |
| Chromogranin A | Abcam | 15160 | 1:100 (IHC/IF) |
| Synaptophysin | Cell Signaling Technologies | 9020 | 1:200 (IHC/IF) |
| Irx-2 | Abcam | 196755 | 1:100 (IHC/IF) |
| Ki-67 | Cell Signaling Technologies | 9449 | 1:400 (IHC/IF) |
| α-smooth muscle actin | Abcam | 5694 | 1:200 (IHC/IF) |
| Glucagon | Cell Signaling Technologies | 2760 | 1:100 (IHC/IF) |
| Insulin | Cell Signaling Technologies | 3012 | 1:1000 (IHC/IF) |
| GFAP (Rabbit pAb) | Dako Agilent Technologies | Z033429 | 1:400 (IHC/IF)<br>1:2,000 (WB) |
| GFAP | Invitrogen | PA1-10004 | 1:400 (IHC/IF) |
| Nestin | Novus Biologicals | NB100-1604 | 1:500 (IHC/IF)<br>1:100 (ICC/IF) |
| Sox2 | Novus Biologicals | NB110-37235 | 1:200 (ICC/IF) |
| Pit-1 | Novus Biologicals | NBP1-92273 | 1:500 (IHC/IF) |
| Prolactin | Abcam | 183967 | 1:200 (IHC/ICC/IF) |
| Gastrin | Dako Agilent Technologies | A0568 | 1:1000 (IHC/IF) |
| Acetylated-tubulin | Sigma | MABT868 | 1:200 (IHC/IF) |
| Gastrin-releasing peptide | Invitrogen | PA5-115272 | 1:200 (IHC/IF) |
| GAPDH | Cell Signaling Technologies | 5174S | 1:2,000 (WB) |
| Gli2 | Santa Cruz Technologies | 271786 | 1:1,000 (WB) |
| Menin (Mouse mAb) | Bethyl Laboratories | A500-003A | 1:100 (IP) |
| GFAP (Rat mAb) | Invitrogen | 13-0300 | 1:100 (IP) |

## ABBREVIATIONS

BG: Brunner’s glands
ChgA: Chromogranin A
Gcg: Glucagon
GEP-NENs: gastroenteropancreatic neuroendocrine neoplasms
GFAP: Glial fibrillary acidic protein
Grp: Gastrin-releasing peptide
Ins1: Insulin
MEN1: Multiple endocrine neoplasia 1
nPan: Normal pancreas
nPit: Normal pituitary
PCA: Principal Component Analysis
PitNET: Pituitary neuroendocrine tumor
PNEC: Pancreatic neuroendocrine carcinoma
PNET: Pancreatic neuroendocrine tumor
Prl: Prolactin
SMA: Smooth muscle actin
siRNA: small interfering RNA
Sst: Somatostatin
Syp: Synaptophysin
Vip: Vasoactive intestinal peptide

## FIGURE LEGENDS

**Table 1. Summary of phenotypes observed in *GFAP Cre; Men1* mice aged 13-24 months**.

**Supplementary Figure 1.**
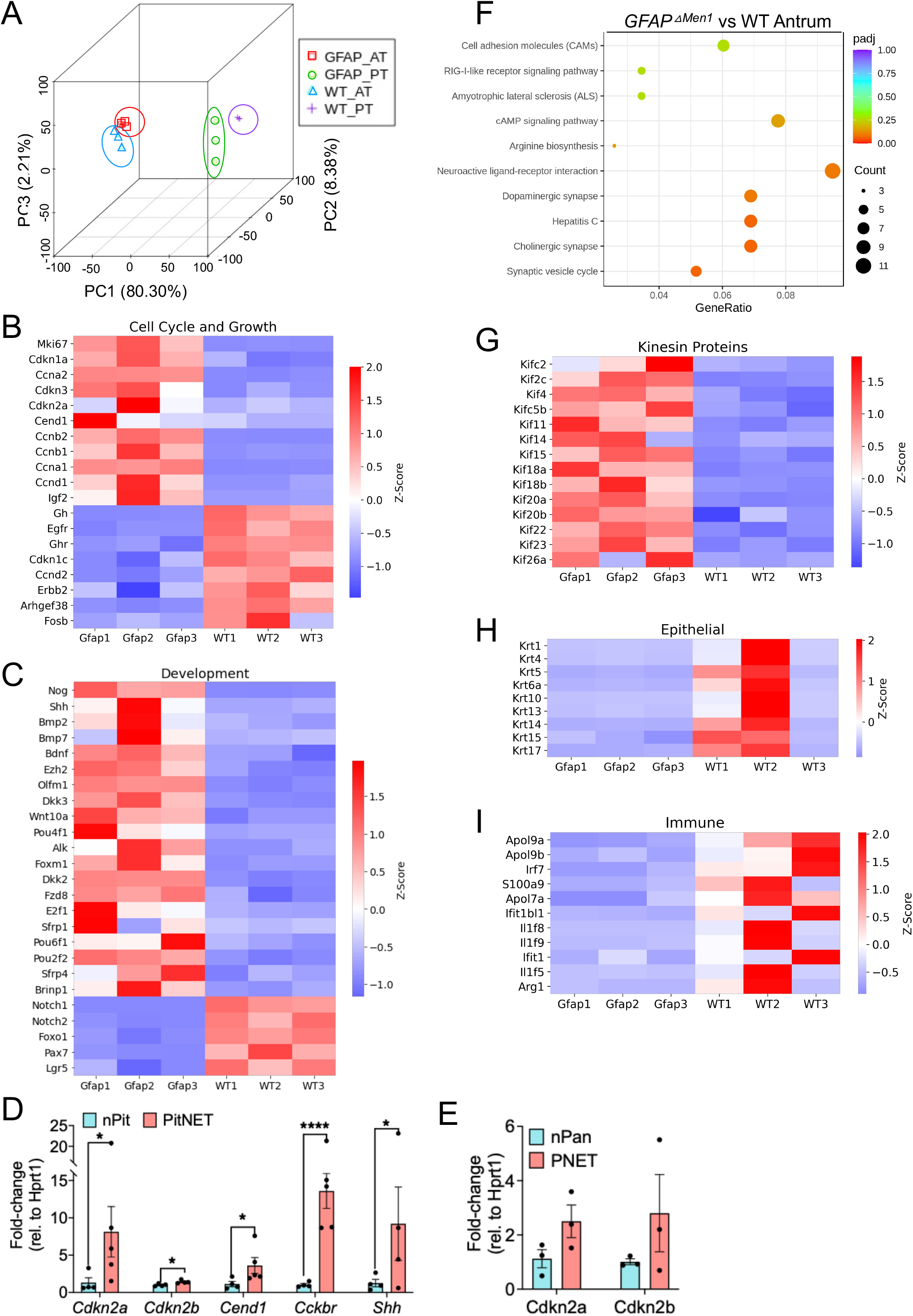
(A) Principal Component Analysis plot capturing transcriptome-wide variation in three components. (B) Heatmaps of significant DEGs mapped to cell cycle regulation and (C) development in *GFAP*^*△Men1*^ pitNETs. (D) Validation of select DEGs in pitNETs versus wild type pituitary by quantitative PCR. n=4–5 mice per group. * *p* < 0.05, *** *p* < 0.001 by Two-way ANOVA. Data are represented as mean ± SEM. (E) Cross validation of select pitNET-enriched transcripts in *GFAP*^*△Men1*^ PNETs and wild type pancreas. (F) KEGG Ontology Pathway analysis of antra from wild type and *GFAP*^*△Men1*^ mice showing the number of genes mapped to enriched pathways and their level of statistical significance. (G) Heatmap showing enrichment of kinesin motor protein transcripts in *GFAP*^*△Men1*^ pitNETs. (H) Heatmaps depicting downregulation of cytokeratins and (I) immune-related transcripts in *GFAP*^*△Men1*^ antral extracts. n=4–5 mice per group with the exception of wild type pituitary group which represents four samples of three pooled pituitaries for a total of 12 tissues in this group.

